# Computational verification of large logical models – application to the prediction of T cell response to checkpoint inhibitors

**DOI:** 10.1101/2020.05.01.073379

**Authors:** Céline Hernandez, Morgane Thomas-Chollier, Aurélien Naldi, Denis Thieffry

## Abstract

At the crossroad between biology and mathematical modelling, computational systems biology can contribute to a mechanistic understanding of high-level biological phenomenon. But as knowledge accumulates, the size and complexity of mathematical models increase, calling for the development of efficient dynamical analysis methods. Here, we propose the use of two approaches for the development and analysis of complex cellular network models.

A first approach, called “model verification” and inspired by unitary testing in software development, enables the formalisation and automated verification of validation criteria for whole models or selected sub-parts. When combined with efficient analysis methods, this approach is suitable for continuous testing, thereby greatly facilitating model development.

A second approach, called “value propagation”, enables efficient analytical computation of the impact of specific environmental or genetic conditions on the dynamical behaviour of some models.

We apply these two approaches to the delineation and the analysis of a comprehensive model for T cell activation, taking into account CTLA4 and PD-1 checkpoint inhibitory pathways. While model verification greatly eases the delineation of logical rules complying with a set of dynamical specifications, propagation provides interesting insights into the different potential of CTLA4 and PD-1 immunotherapies.

Both methods are implemented and made available in the all-inclusive CoLoMoTo Docker image, while the different steps of the model analysis are fully reported in two companion interactive jupyter notebooks, thereby ensuring the reproduction of our results.

## 1 INTRODUCTION

Recent technical developments have allowed scientists to study immunology and health-related issues from a variety of angles. For many diseases, especially for cancer, the current trend consists in aggregating data coming from different sources to gain a global view of cell, tissue, or organ dysfunction. Over the last decades, diverse mathematical frameworks have been proposed to seize a multiplicity of biological questions (Le Novère, 2015), including in immunology (Kaufman et al., 1985, 1999; Eftimie et al., 2016; Chakraborty, 2017). However, the increasing complexity of biological questions implies the development of more sophisticated models, which in turn bring serious computational challenges.

Among the mathematical approaches proposed for the modelling of cellular networks, the logical modelling framework is increasingly used. In particular, it has been successfully applied to immunology and cancer, leading to the creation of models encompassing dozens of components, some including many inputs components (Grieco et al., 2013; Abou-Jaoudé et al., 2014; Flobak et al., 2015; Oyeyemi et al., 2015). However, the large size of recent models hinders the complete exploration of their dynamical behaviour through simulation, especially in non-deterministic settings.

To address these difficulties, we define and apply a *model verification* approach to systematically verify whether a model complies with a list of known properties (section 2). These properties are defined as model *specifications*, either at a local (i.e. for sub-models) or at a global level. This automated verification procedure fosters confidence during the development of a complex dynamical model and paves the way to the development of models with hundreds of nodes.

We further outline and apply a *value propagation* method, which enables the assessment of the impact of environmental or genetic constraints on the dynamical behaviour of complex cellular networks (section 3).

These two complementary approaches can be applied to the development and analysis of large dynamical models, as illustrated in Figure 1. Noteworthy, they have been implemented in a multi-platform Docker image combining various complementary logical modelling and analysis tools (Naldi et al., 2018b). We further illustrate the power of these methods through the analysis of an original model described in Section 4. The different steps of analysis are fully reported in two companion interactive jupyter notebooks, available with the model on the GINsim website (http://ginsim.org/model/tcell-checkpoint-inhibitors-tcla4-pd1), thereby ensuring their reproducibility.

**Figure 1.**
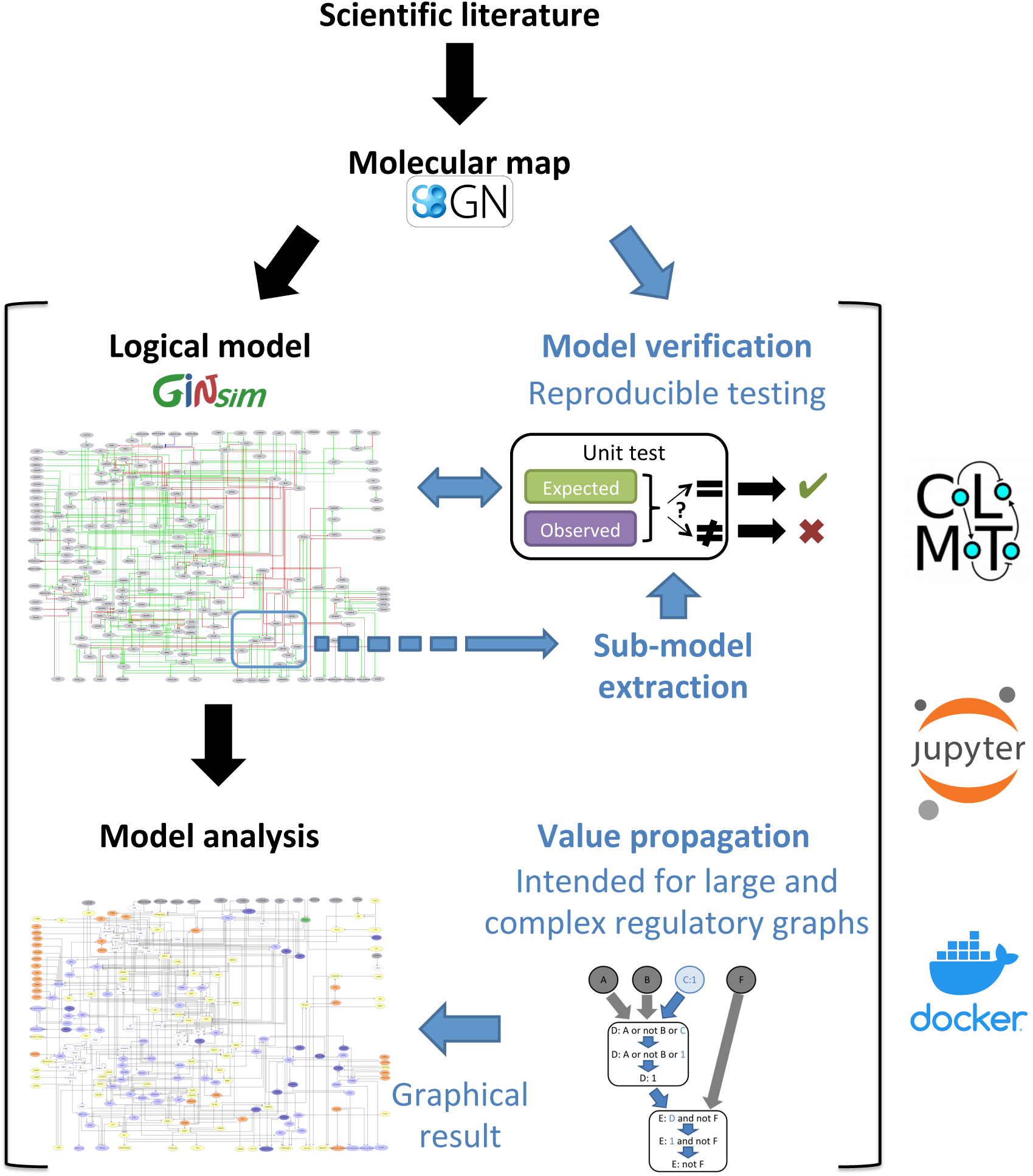
Description of the proposed workflow for the development and analysis of dynamical logical models. The novel methods described in this article are emphasized with blue fonts. Starting with the delineation of a molecular map integrating the available scientific knowledge, we derive a regulatory graph and logical rules to generate a logical model, and induce dynamical specifications serving as test cases to verify the model. Moreover, when the available knowledge is specific to a smaller part of the regulatory graph, a sub-model us extracted to perform local tests. We further implemented an analysis and visualisation method, called Value propagation, to assess the impact of environmental and genetic perturbations. Figure 3 zooms into this part of the workflow and describes it in more details. The use of model verification, sub-model extraction and value propagation is illustrated in two reproducible and editable Jupyter notebooks, taking advantage of the CoLoMoTo Interactive notebook framework (Naldi et al., 2018b). This framework is available inside the CoLoMoTo Docker image together with packaged libraries for the analysis of dynamical logical models of biological networks.

## 2 MODEL VERIFICATION

### 2.1 A software engineering framework for logical model building

One of the main features determining the interest of a model is its ability to accurately recapitulate salient biological knowledge. More precisely, this knowledge can be used in two complementary ways during the model building process. On the one hand, it is used to define the model architecture, specifying which biological entities need to be included and which interactions between these entities need to be encoded. On the other hand, biological knowledge entails dynamical properties that must be achieved by the resulting model, whether transitory or asymptotic, to account for biological observations. These properties induce satisfaction criteria and must be clearly specified for rigorous model assessment or comparison with other models. Failures to reproduce such properties need to be carefully documented, thereby providing a basis for further model improvement.

In the domain of logical modeling applied to cellular networks, various formal methods have already been proposed to verify dynamical properties. For example *stable states* (or textitfixed points, characterised by all components being steady at the same time) tentatively correspond to asymptotic properties that can used to assess the reproduction of known persistent biological behaviour. More complex asymptotic behaviours include *cyclic attractors*, which can be approximate by the computation of so called *trap spaces*. Also called *stable motifs*, trap spaces are hypercubes in the state space such that all successors of all states in the hypercube also belong to it (for synchronous and asynchronous updatings, or any other updating). These hypercubes then provide an approximation of complex attractors. Trap spaces and stable states can be defined as results of a constraint solving system, enabling their efficient computation (Klarner et al., 2018). Their reachability however must be assessed separately, often using model checking or stochastic simulations, which requires longer computations.

Model checking techniques have been successfully applied to specify and verify temporal constraints on a model behaviour (Monteiro and Chaouiya, 2012; Miskov-Zivanov et al., 2016; Traynard et al., 2016; Wang et al., 2016).

In any case, whatever the formalism chosen, the building of a complex dynamical model is intrinsically iterative, as its establishment is usually incremental and requires continuous testing and adjustment with reference to a growing body of biological knowledge.

In the field of software engineering, the similar need to repeatedly assess criteria of success or failure of a software program led to the development of powerful *software verification* techniques, and in particular to software testing (Myers, 1979), which main goal is to assess whether a software meets a series of well-defined requirements. More importantly, such assessments must be repeated as soon as a new piece of code or specification is added. Software testing aims to check whether newly introduced modification might break any of the previous performances. In particular, software verification includes the notion of *unit testing*, where suites of tests describe the expected behaviour associated with individual units composing a program. This idea can be transposed from computer science to model building and has been successfully applied in the context of other modeling frameworks (Hoops et al., 2006; Lopez et al., 2013; Sarma et al., 2016; Boutillier et al., 2018), but not yet to logical modeling.

Here, we transpose the *unit testing* approach to integrate a comprehensive series of verifiable criteria, from the early stages of model conception, in order to fully automate the dynamical evaluation of logical models. The core idea is to split the biological knowledge on which a model is based into individual verifiable criteria that can be formalised as specifications (Figures 1 and 2). In this respect, individual units of knowledge, derived from the scientific literature or biological experiments, must be formulated into stable or dynamical properties. Each specification, coupling a property with an expected value, can serve as a basis to define a test case for a model. Testing such a specification amounts to compute an “observed” value based on the model and compare this value to the expected one.

**Figure 2.**
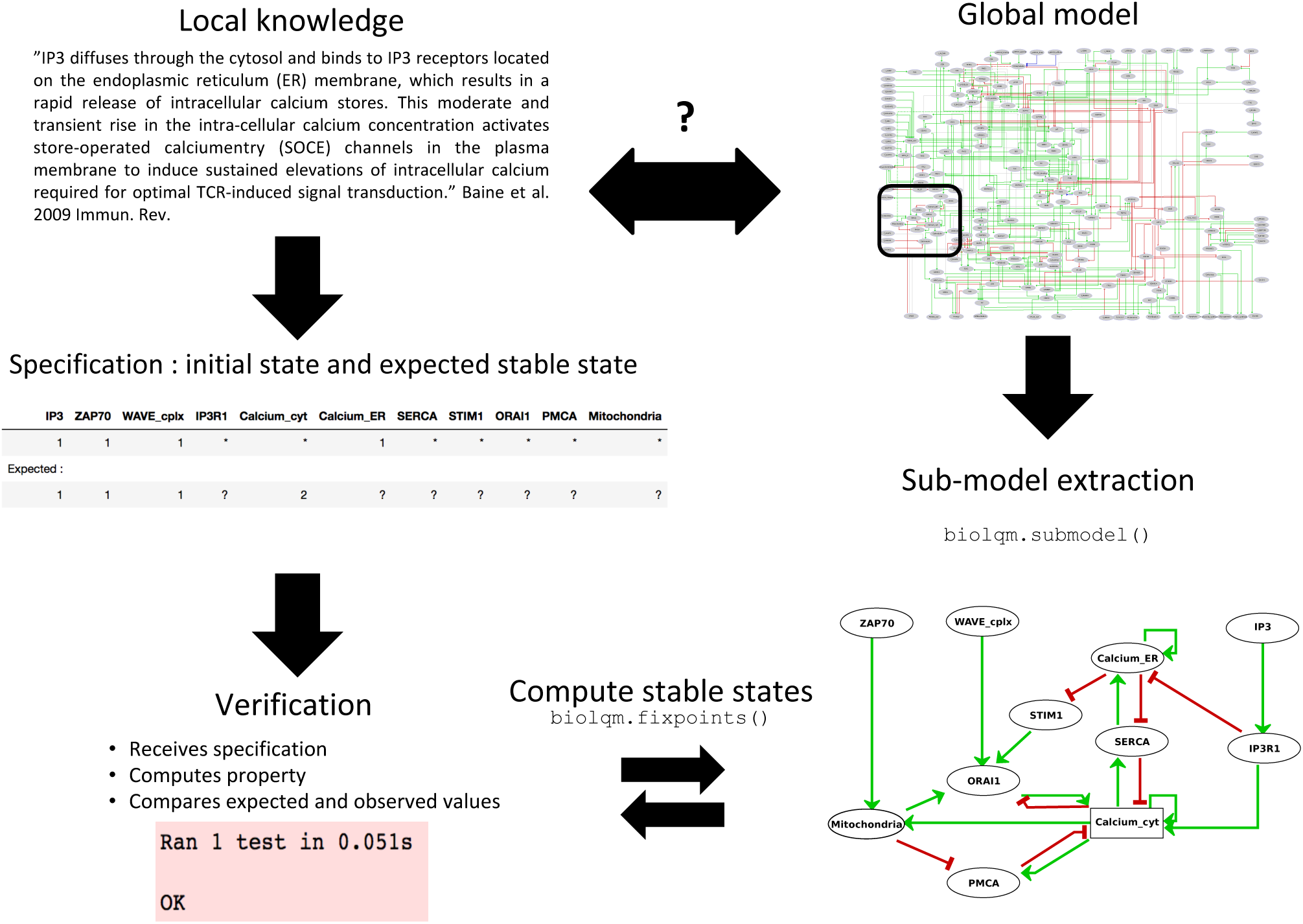
Sub-model extraction for local model verification. When the available knowledge is fragmentary and covers the behaviour of only a subset of components, verification becomes difficult at the global scale of the model. Based on this partial information, a series of specifications can still be defined for a sub-model which, after extraction using bioLQM’s submodel() function, can then be rigorously tested.

In practice, the CoLoMoTo notebook environment (Naldi et al., 2018b) provides a Python API for several software tools, enabling the definition of a wide range of dynamical analysis for the computation of observed values. Individual test cases can be assembled into a library, also called *testing suite*. Existing tools and packages enabling software testing can then be applied to automatically assess whether a model satisfies (or not) a series of specifications. In this study, we used the python package ‘unittest’, taking advantage of its seamless integration into the CoLoMoTo interactive notebook. This unit-testing package is integrated by default into the recent versions of the Python standard library (http://python.org).

### 2.2 Local verification of sub-models can cope with sparsity of biological knowledge

Biological knowledge reported in the scientific literature is often insufficient to evaluate a comprehensive model, which may encompass hundreds of nodes. In particular, observations regarding component activity often relate to only a limited subset of nodes of the model. This greatly complicates the definition of specifications for the whole model.

Given a comprehensive model and a set of components of interest, one can extract a sub-model containing these core components, along with their associated logical rules. Components which take part of these logical rules but are not part of the selected set are considered as external inputs of the sub-model (Figure 2). This functionality has been implemented in the ‘submodel’ function of the Java bioLQM library (Naldi, 2018) according to the following procedure.

Let *M* = (*V, f*) be a model, where *V* is the set of components, and *f* the update function. For each *c* in *V, f*_*c*_ is the logical function associated to the component *c* and *R*(*c*) is the set of its regulators (i.e. components that intervene in the logical rule). Given a list of selected components *C ⊂ V* :

1. *S* = Ø
2. for each component *c* ∈ *C*: *S* = *S* + {*c*} + *R*(*c*)
3. create the sub-model *M*′ = (*S*; *f*′) such that for each component *c* in 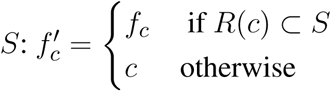

As shown in Section 4, the delineation of such sub-models can greatly facilitate the definition and verification of local specifications.

## 3 VALUE PROPAGATION ENABLES THE EVALUATION OF THE IMPACT OF A GIVEN CELLULAR ENVIRONMENT ON MODEL DYNAMICS

The core idea of *value propagation* is presented in Figure 3. Given a set of logical rules and a cellular context, an iterative algorithm enables the computation of the dynamical consequences of the cellular context on all the components of the model.

**Figure 3.**
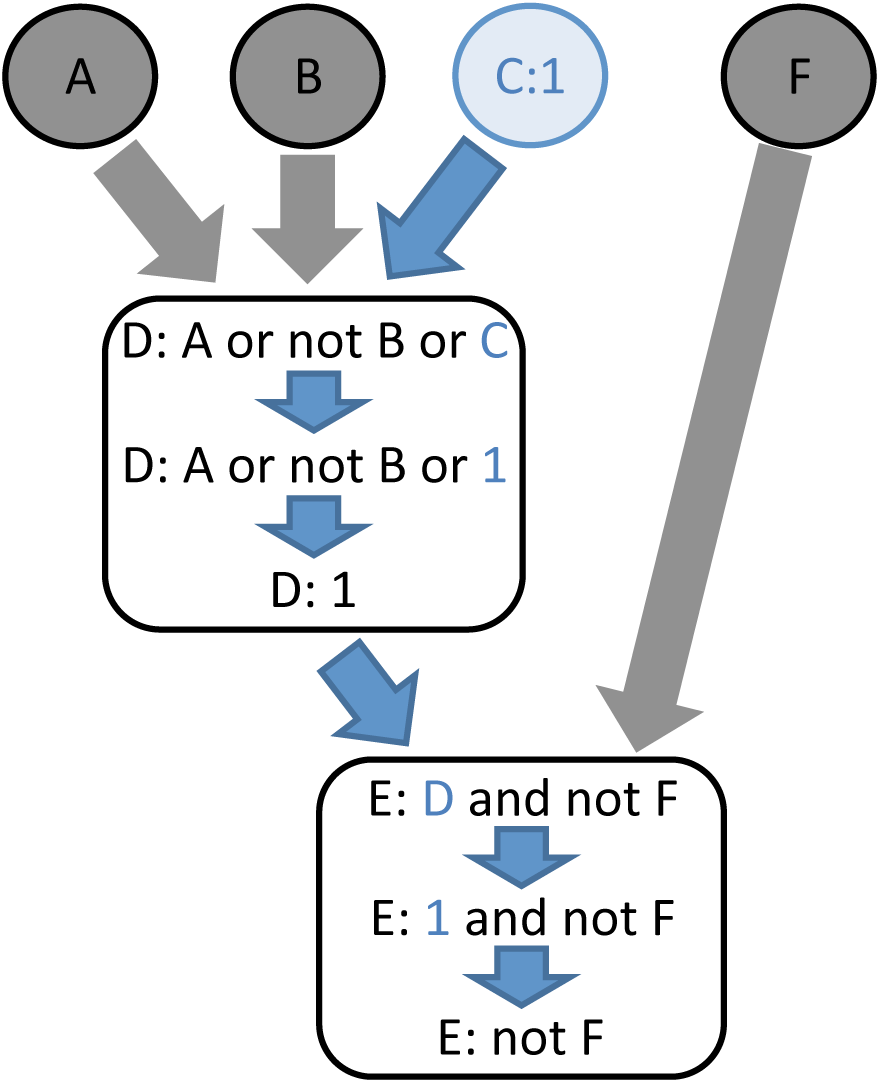
The principle of logical value propagation analysis is illustrated with a simple example involving two core nodes, D and E, and four input nodes, A, B, C and F. The value 1 is assigned to the node C and then propagated through the model. The assignment C:1 implies the evaluation of D to 1. Consequently, the function assigned to node E becomes ‘not F’. In other words, assigning the value 1 to node C activates node D independently from the value of its other inputs, while node E becomes completely dependent on the value of node F.

First, the cellular context is formalised by assigning constant values to some components of the model. Next, we apply a recent model reduction technique reported by Saadatpour *et al*. (Saadatpour et al., 2013). Briefly, for each constant node, the corresponding value is inserted into the logical rule associated with each of its target nodes. Each logical rule is then simplified using Boolean algebra. If the rule simplifies to a constant, this fixed value is further propagated into the logical rules of downstream nodes. This process is iterated until no further propagation or simplification can be made on the logical rules of the model. In contrast with the approach of Saadatpopour *et al*., which aims at producing a reduced model, we focus principally on the outcome of the propagation of fixed values.

The result of value propagation can be very informative by itself. Indeed, the resulting stabilised values provide insights into the impact of a given (single or multiple) perturbation on the model, revealing which elements are consequently constrained to become activated or inactivated, versus which elements keep some degree of freedom. Furthermore, this method greatly eases the comparison of the impacts of different biological contexts on network dynamics by performing a differential analysis of the corresponding lists and target values of fixed components. This method has multiple advantages when applied to complex networks, as it can be used efficiently on models with large numbers of components. It further simplifies the computation of attractors (stable states or even simple or complex cycles). Interestingly, Saadatpour and collaborators showed that this method conserves the stable states and complex attractors under the fully asynchronous updating assumption (Saadatpour et al., 2013).

This method was extended to multilevel models and implemented into the Java bioLQM library (Naldi, 2018). In this implementation, the fixed components are conserved during value propagation, enabling a direct comparison of the propagated effects of alternative perturbations.

The power of this approach is demonstrated on a concrete example in the following section.

## 4 APPLICATION: ASSESSING THE EFFECT OF CHECKPOINT BLOCKADE THERAPIES ON T CELL ACTIVATION

### 4.1 Biological background

Over the last decades, immunotherapies have been the subject of intense studies and led to great advances in the field of cancer treatment. Through the years, it has then been recognised that T cells often display a reduced ability to eliminate cancer cells, and that expression of co-inhibitory receptors at their surface accounts for this compromised function. Receptors like Cytotoxic T-lymphocyte protein 4 (CTLA4, also known as CD152) (Walunas et al., 1994; Leach et al., 1996) and Programmed cell death protein 1 (PD-1, also known as PDCD1 or CD279) (Ishida et al., 1992) have been particularly studied in that context. Antibodies blocking the pathways downstream of these co-inhibitors (checkpoint blockade therapies) have become standard treatment for metastatic melanoma (Robert et al., 2011; Simpson et al., 2013) and other cancers (Ribas and Wolchok, 2018), including non-small cell lung cancer, renal cell carcinoma, Hodgkin’s lymphoma, Merkel cell carcinoma and many others. The successes of these studies led to an increasing interest in T cell co-inhibitory receptors.

Nevertheless, a clear understanding of the mechanisms at work inside T cells remains elusive. Therapies targeting CTLA4 or PD-1 show different immune adverse effects (June et al., 2017), while the corresponding intra-cellular mechanisms remain to be clarified. Moreover, a rationale for the educated development of new immunotherapies focusing on other receptors or combinations of receptors is clearly needed. Co-inhibitory receptors are legions at the surface of T cells (Brownlie and Zamoyska, 2013) and biology of T cell activation or tolerance involves activation or repression of highly interconnected and complicated pathways (Baumeister et al., 2016).

Given the central role of T cells in many medical contexts, several mathematical frameworks have been applied to model T cell activation. Recent examples include rule-based approaches (Chylek et al., 2014), ordinary differential equations (Perley et al., 2014), and logical models (Oyeyemi et al., 2015; Rodríguez-Jorge et al., 2019; Sánchez-Villanueva et al., 2019), considering different biomedical contexts as diverse as HIV infection or neonate vaccination. To our knowledge, none of them specifically focused on the impact of co-inhibitory receptors on T cell activation or tolerance.

In this study, we applied the logical framework to integrate current data on CTLA4 and PD-1 pathways and assess their impact on T cell activation. Our goal was triple. First, we wanted to create a comprehensive model building upon extensive knowledge encoded into a molecular map (see next section). Second, using model verification and a specific unit test suite, we aimed to firmly anchor the model at both the global and local scale into the collected biological knowledge. Third, using value propagation, we aimed to provide a tool for the comparative analysis of intra-cellular consequences when targeting CTLA4 versus PD-1 T-cell co-receptors.

### 4.2 Comprehensive molecular mapping of T Cell activation network

Prior to mathematical modelling, knowledge about biological entities involved in T Cell activation was collected from available pathway databases, including Reactome (Fabregat et al., 2016), PantherDB (Mi et al., 2013), ACSN (Kuperstein et al., 2015) and WikiPathways (Slenter et al., 2018). Moreover, the scientific literature indexed in the PubMed database was further explored and carefully curated. Using the software CellDesigner (version 4.3.1) (Funahashi et al., 2008), this knowledge was encoded in a molecular map describing reactions between biological entities (either proteins, RNAs, genes, complexes or metabolites). Each biological entity included in the map was annotated with a series of standard identifiers, including UniProtKB accession number, recommended and alternative names, gene name and synonyms, and cross-references to unique HGNC identifiers and approved symbols. The annotations also reference relevant scientific articles, including PubMed identifier, first and last authors, year of publication, and a list of observations extracted from these publications.

Our T cell activation map currently encompasses 726 biological entities, in different states (active/inactive, with or without post-translational modifications), and 539 reactions involving these entities (Supplementary Figure 1 and File 1). Globally, the map currently integrates information from 123 scientific articles, which are cited in the annotations of the entities and reactions of the map.

### 4.3 Logical modelling of T Cell activation

Using the logical modelling software *GINsim* (version 3.0.0b) (Naldi et al., 2018a), we then manually derived a regulatory graph encompassing 216 nodes and 451 arcs (Figure 4) from the content of the molecular map. One by one, biological entities represented in the molecular map were re-created as components of the logical model. In most of the cases, the representation of entities having different states was further compressed into a single component summarizing their activity in the TCR signalling cascade. Furthermore, to obtain a dynamical logical model, a specific logical rule must be assigned to each node. In many cases, this can be achieved rather easily based on published data. For more complex situations, a default generic logical rule was initially considered, where all activators are needed for the activation of a component (using the AND operator) and where only one inhibitor is sufficient to repress it (using the OR and NOT operators), which served as a basis for further rule refinement. In some cases, however, in particular when a component is the target of various regulatory interactions or when metabolites are involved, finding direct support for a specific rule may be tricky or impossible. Hence, the delineation of consistent logical rules for a complex model is often the result of an iterative process, starting with generic rules and progressively correcting them based on the results of various analyses.

**Figure 4.**
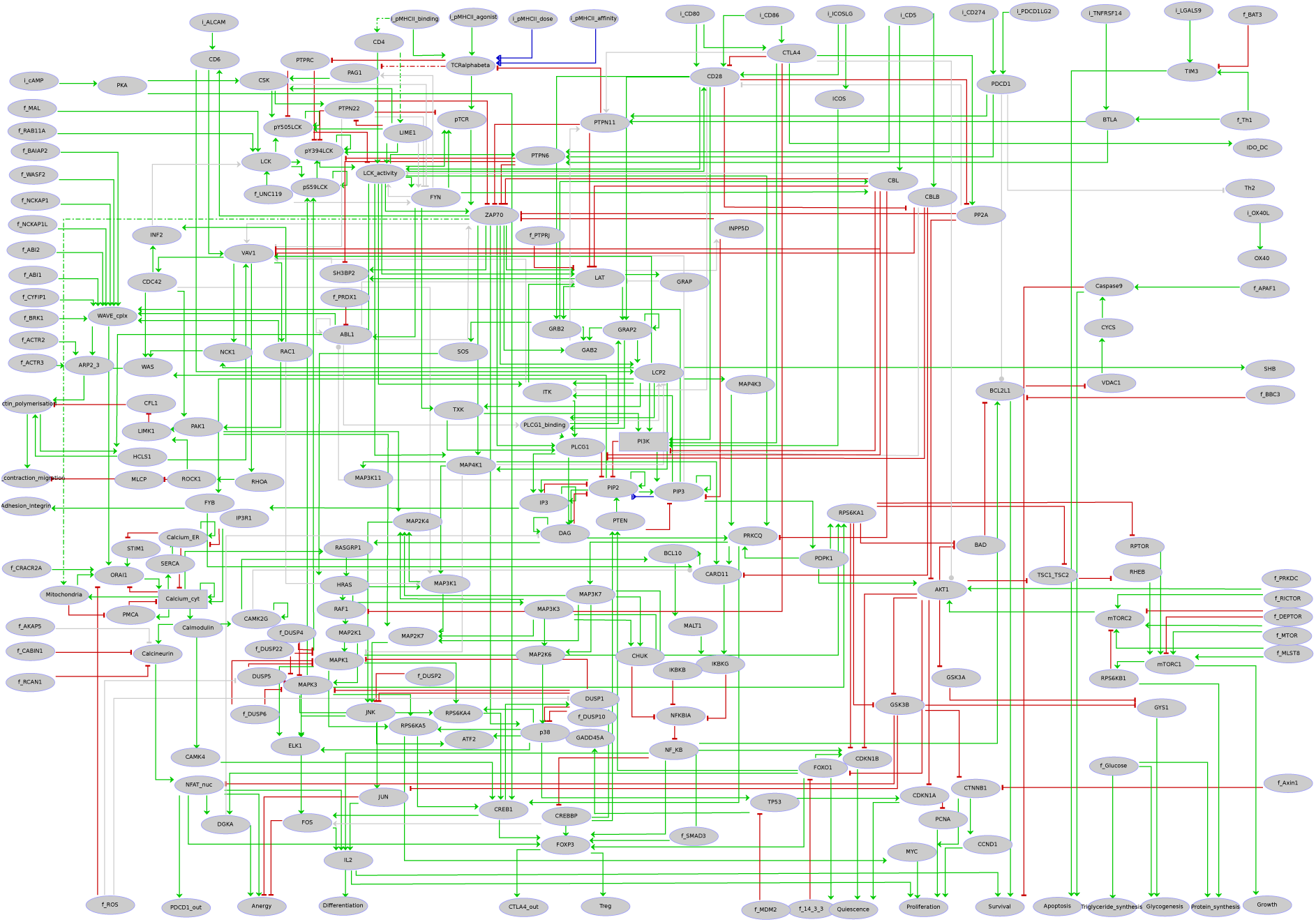
Regulatory graph of the T cell activation model. The global layout is similar to the molecular map (cf. supplementary Figure 1 and supplementary File 1), with ligands/receptors and proximal signalling at the top, and the nucleus-related events at the bottom of the graph. In between, the model encompasses interconnected pathways and signalling cascades related to cytoskeleton remodelling, the MAPK network, calcium fluxes, metabolic shifts, and NF-*κ*B, to name a few. Boolean components are denoted by ellipsoids whereas rectangles denote ternary components. Green arcs denote activation events, red blunt arcs denotes inhibitions, while blue arcs denote dual regulations. The grey arcs represent interactions created during the translation of the molecular map into the regulatory graph, but that are not yet integrated at the dynamical level (i.e. not taken into account in the logical rule).

Hereafter, we demonstrate how we can take advantage of the methods presented in the previous sections to ease rule refinement by model verification. We first defined a series of properties expected for the model (see examples in Table 1). Next, stable states and/or trap spaces were computed and automatically compared with these properties (cf. first Jupyter notebook provided on the model web page at http://ginsim.org/model/tcell-checkpoint-inhibitors-tcla4-pd1). After some iterative runs of the notebook, manual refinements lead us to a set of rules complying with all the tests.

**Table 1.**
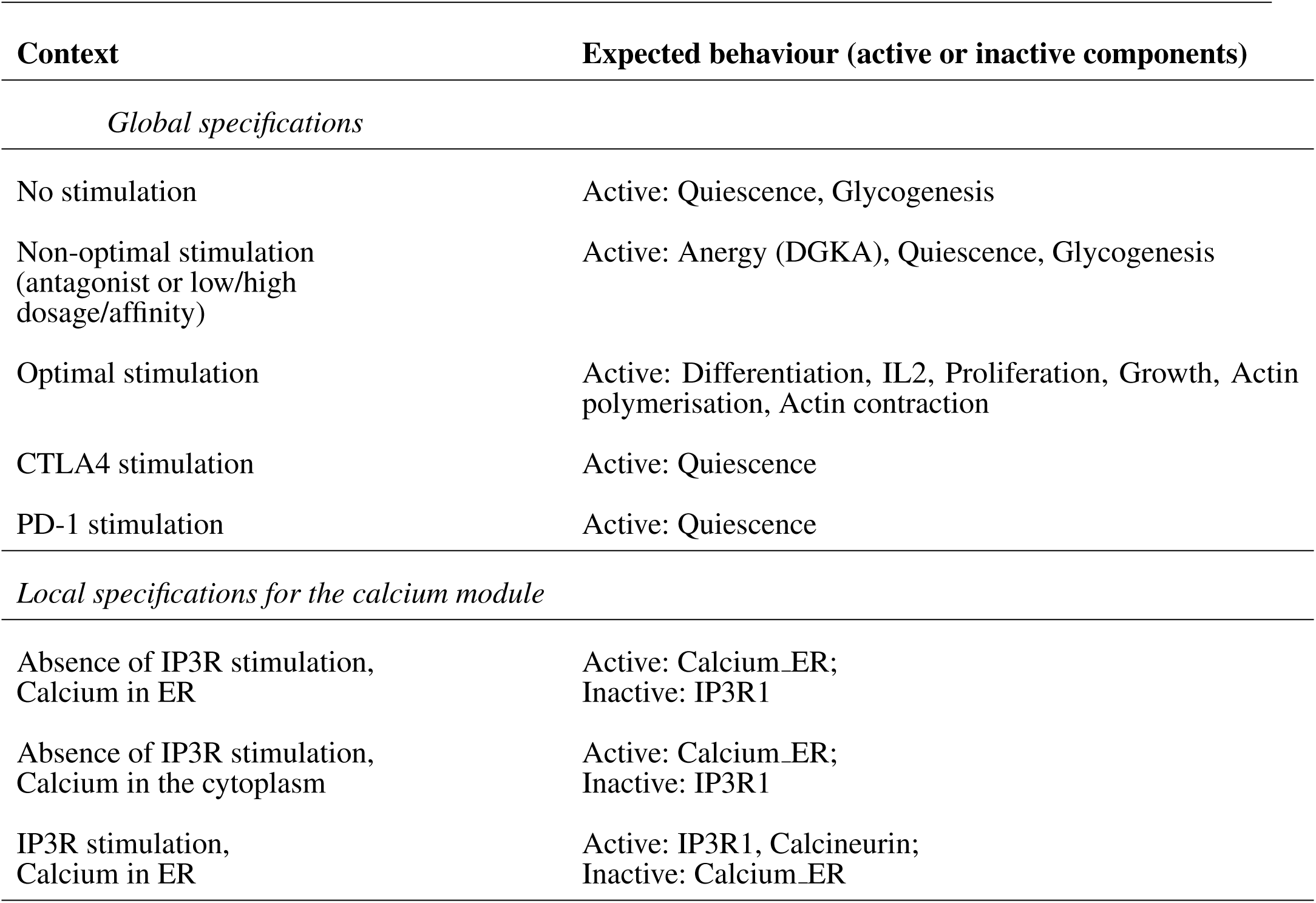
Global specifications used to assess the T cell activation model and example of local specifications for the calcium signaling module (cf Figure 2). After verification, named components should have a value of 0 if specified as inactive, while active components should have a value of 1 or 2. Verification of local specifications requires the extraction of a sub-model from the global model. These verifications and literature references are detailed in the companion CoLoMoTo notebooks. ER: Endoplasmic Reticulum.

| Context | Expected behaviour (active or inactive components) |
| --- | --- |
| *Global specifications* |  |
| No stimulation | Active: Quiescence, Glycogenesis |
| Non-optimal stimulation<br>(antagonist or low/high dosage/affinity) | Active: Anergy (DGKA), Quiescence, Glycogenesis |
| Optimal stimulation | Active: Differentiation, IL2, Proliferation, Growth, Actin polymerisation, Actin contraction |
| CTLA4 stimulation | Active: Quiescence |
| PD-1 stimulation | Active: Quiescence |
| *Local specifications for the calcium module* |  |
| Absence of IP3R stimulation,<br>Calcium in ER | Active: Calcium_ER;<br>Inactive: IP3R1 |
| Absence of IP3R stimulation,<br>Calcium in the cytoplasm | Active: Calcium_ER;<br>Inactive: IP3R1 |
| IP3R stimulation,<br>Calcium in ER | Active: IP3R1, Calcineurin;<br>Inactive: Calcium_ER |

For example, the Endoplasmic Reticulum (ER) serves as a reservoir for calcium ions. This reservoir can be emptied through activation of the Inositol 1,4,5-trisphosphate receptor (IP3R1). When empty, this reservoir can be filled through activation of the Sarcoplasmic/endoplasmic reticulum calcium ATPase 2 (SERCA) pumps. A default logical rule for a node representing the presence of this Calcium quantity (Calcium ER) is then ‘SERCA AND NOT IP3R1’. To check the behaviour of the corresponding logical sub-model, we defined a test checking whether whenever Calcium_ER was evaluated to TRUE, SERCA was evaluated to FALSE (see test ‘test_calc_tp_rest_ER1_SERCA0’). However, consecutive model verification failed, allowing us to notice that the default rule implied that SERCA should be always TRUE for Calcium_ER to be TRUE. The rule was then corrected to take into account the fact that Calcium_ER should stay TRUE whenever it would reach this value in absence of IP3R1.

In the first Jupyter notebook provided as supplementary material, we include all the code enabling the verification of our final model, which encompasses 36 unit tests split in four test suites. On a MacBook Pro using macOS 10.13 High Sierra, with a 2.3 GHz Intel Core i7 and 16GB 1600 MHz DDR3, all the tests were run in 87s.

The four test suites cover the most complex parts of the model, some of them particularly difficult to define. These suites use sub-models, whose delineation was guided by known pathways and practical knowledge gained by the modeler during the assembly of the molecular map. The Calcium module test suite covers a sub-model related to the fluxes of Calcium ions between different cellular compartments, namely the endoplasmic reticulum, the cytoplasm and the extracellular region. The LCK module test suite is centered on the Tyrosine-protein kinase Lck (LCK). This kinase is known to have multiple sites of phosphorylation, whose collective status determines the tridimensional conformation and thus the activity of the enzyme (Ventimiglia and Alonso, 2013). The Cytoskeleton module test suite covers the cytoskeleton remodelling events occuring during T cell activation, and has strong connections with the Calcium sub-model. Finally, the Anergy/activation/differentiation module covers a less documented module encompassing the nucleus compartment and gene transcription.

### 4.4 Comparison of the impacts of CTLA4 and PD-1 co-inhibitory receptors through value percolation

Based on the model described in the preceding section, a comparative propagation analysis was performed to visualise the respective effects of CTLA4 and PD-1 receptor activation on model dynamics. Figure 5 displays the value propagation for each condition on a single regulatory graph, using a color code to distinguish the different situations (component inhibition/activation in one or both conditions). The value propagation for the two conditions are further shown separately in the second companion notebook (available at http://ginsim.org/model/tcell-checkpoint-inhibitors-tcla4-pd1). This analysis reveals that the activation of the CTLA4 receptor impacts most pathways of the model, impeding in particular the remodeling of the cytoskeleton and the metabolic switch associated with bona fide T cell activation. In contrast, the activation of the PD-1 receptor leads to more limited effects, predominantly freezing the components of the NF-*κ*B pathway.

**Figure 5.**
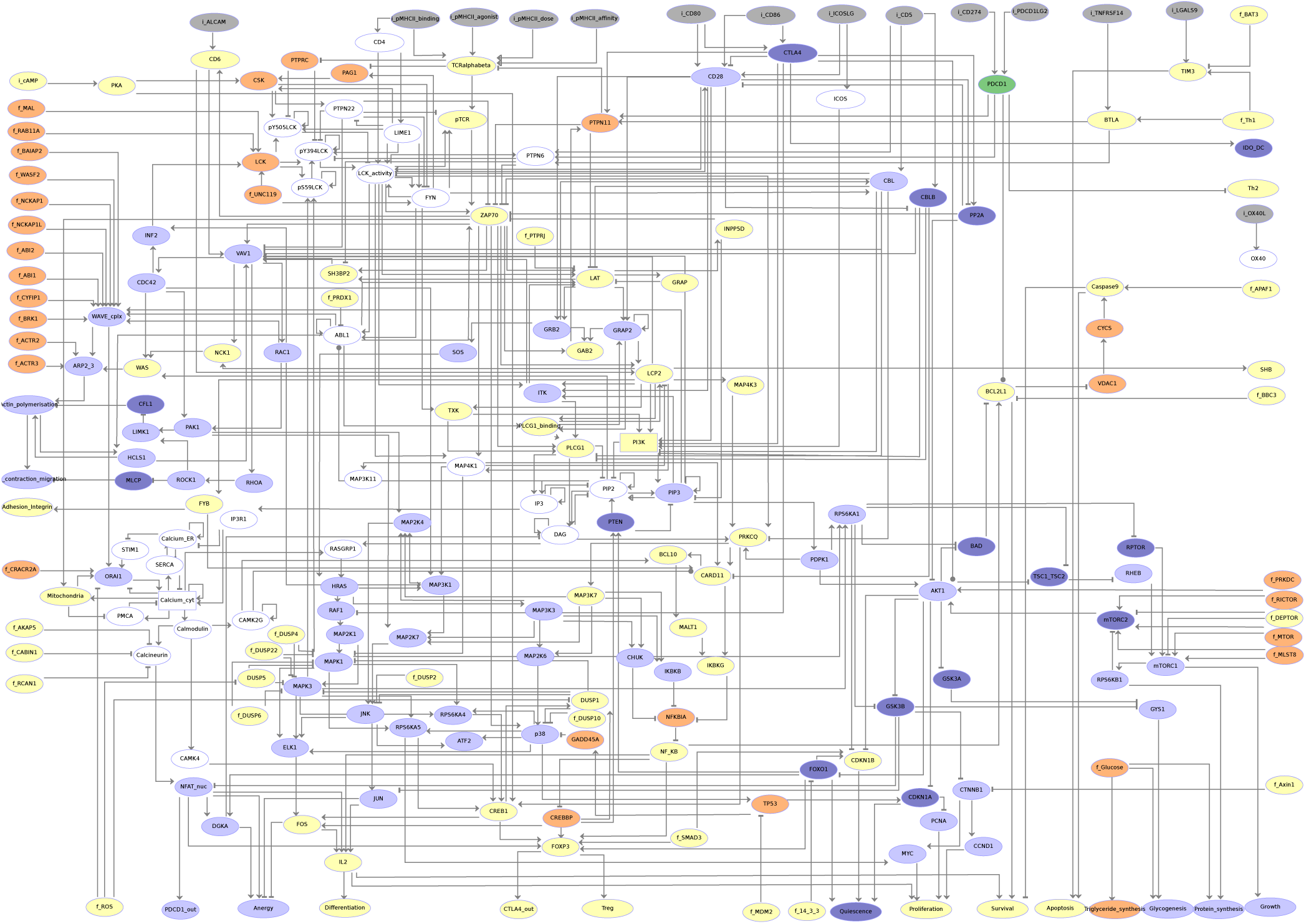
Visualisation of the results of the propagation analyses for CTLA4 versus PD-1 activation. Grey nodes correspond to inputs. Nodes in yellow are frozen OFF upon any of CTLA4 or PD-1 (PDCD1) activation. Nodes in orange are frozen ON (i.e. with level 1 or 2) for each of these conditions. Nodes in light blue are frozen OFF only for CTLA4 activation (i.e. they remain free upon PD-1 activation). Nodes in dark blue are frozen ON (i.e. with level 1 or 2) only upon CTLA4 activation (i.e. they remain free upon PD-1 activation). Upon PD-1 activation, the corresponding node (PDCD1) is the only one that gets specifically frozen (ON, shown in dark green). Nodes in white remain free for both conditions.

A more refined comparative analysis of value propagation from these two receptor activations entails the observation that the set of nodes frozen by the propagation of PD-1 activation is completely included inside the set of nodes frozen by the propagation of CTLA4 activation (see Table 2). Furthermore, the values of the components frozen in both propagation studies are the same. Interestingly, a set of nodes related to calcium influx from and to the endoplasmic reticulum remain unfixed by any of the propagation analyses. This could be an artefact of the positive feedback loops added on the nodes representing the Calcium ion levels in different compartments and would need to be further investigated. A more detailed biological interpretation of these results is proposed in the following section.

**Table 2.**
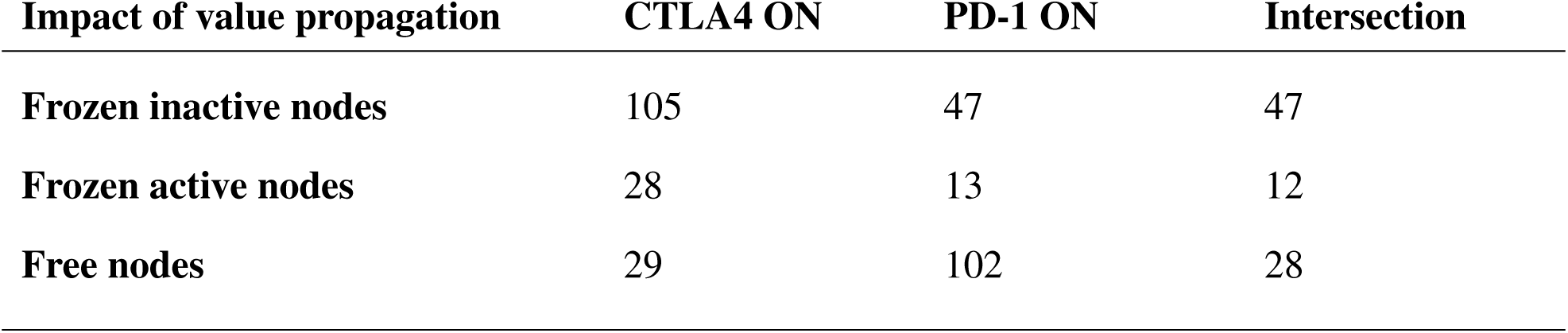
Quantification of the model nodes impacted by the propagation of CTLA4 or PD-1 persistent activation. After propagation, the nodes of the model can remain free (not fixed) or become frozen inactive (value 0) or frozen active (value 1, or potentially higher in the case of multilevel components). The model encompasses a total of 216 nodes, including 14 inputs and 40 nodes not affected by CTLA4 or PD-1 activation. Interestingly, PD-1 itself is the only single node specifically frozen by its activation (and not by the activation of CTLA4). Furthermore, each of 59 nodes affected by the two perturbations is frozen to the same value in both cases. Details of the computation method can be found in the companion CoLoMoTo notebooks.

## 5 CONCLUSIONS AND PROSPECTS

In this study, we have implemented and applied two complementary methods enabling a specification-oriented model building approach, thereby easing the delineation and analysis of highly complex logical models. In this respect, the building of a knowledge base, e.g. in terms of a molecular map, is an important first step. In the molecular map (provided as the supplementary File 1), we have integrated the most relevant biological references available on T cell activation and inhibition pathways.

This map is clearly due to evolve, in particular thanks to the generation and analysis of novel high-throughput data (see e.g. the recent extensive analysis of the TCR signalosome (Voisinne et al., 2019)). But any modification needs to be manually propagated to the dynamical model. To date, methods to derive proper dynamical models from such molecular maps are still in their infancy. In the particular case of the Boolean framework, only one automated approach has been recently proposed (Aghamiri et al., 2020). However, a limitation of this approach is the generation of generic logical rules based on static knowledge. Hence, the methods presented here could be used to advantageously refine these rules, taking into account additional biological knowledge about the behaviour of the system under study.

We used the information gathered in our T cell activation map to build a dynamical logical model encompassing over 200 components and 450 interactions. For such a complex model, defining the logical rules in concordance with biological knowledge is a difficult and error-prone process, usually involving iterative trial simulations, where failures are identified to suggest potential improvements. Hence, listing comprehensive and consistent model specifications is a crucial step for model construction. These specifications can be revised as the modeller deepens his understanding of the biological processes under study. Noteworthy, such systematic testing procures a sense of confidence during the development process.

In the unit tests developed for our model, the definition of sub-models was guided by biological knowledge and pathway definitions, while relying partly on the modeller intuition. This step could be improved by community analyses of the regulatory graph to improve their definition.

Model checking techniques have been previously applied to assess model behaviour through systematic cycles of model refinements (see e.g. Traynard et al. (2016) and reference therein). Model verification, as defined here, is a generalisation of this approach, as it can rely on any available analysis as long as its result can be compared to an expected outcome. In our hands, in the course of model building, the unit testing approach, strongly anchored to available knowledge, proved to be very efficient to assess and improve model consistency with respect to a list of biological specifications, without the need of time-consuming and costly simulations. Implemented in the CoLoMoTo Interactive notebook framework (Naldi et al., 2018b), this approach enabled us to define a model recapitulating the most salient properties observed in response to T cell activation, including quiescence, anergy and differentiation.

The use of model checking techniques could be further extended to assess the sensitivity of model behaviour to the choice of specific logical rules. Such extension is hindered by the exponential increase of the number of possible logical rule, as the number of regulators increases. We would thus need a rationale to explore the space of logical rules. A first step in this direction can be found in Abou-Jaoudé and Monteiro (2019).

The approach presented here could also be improved by taking into account and tracking uncertainty during model conception (Thobe et al., 2018), or yet by taking advantage of computational repairing methods (Gebser et al., 2010) to identify more precisely remaining inconsistencies with biological data. Furthermore, other software engineering techniques, such as *code coverage*, could be borrowed to further improve model building and verification. As code coverage computes how much of a program’s code is covered by unit tests, one could design a method computing the fraction of the components of a model that is effectively covered by specifications.

Value propagation analysis of our large and complex regulatory graph proved to be biologically insightful. Indeed, this straightforward approach enabled us to clearly contrast the respective impacts of CTLA4 and PD-1 on T cell activation in our model, providing some rationale for their differential effects in current therapeutic studies. Indeed, anti-CTLA4 immunotherapies are known for their strong adverse effects related to autoimmunity and immunotoxicity (June et al., 2017). Anti-CTLA4 immunotherapies are currently combined with anti-PD-1 immunotherapy, known for its milder impact on the immune system.

Interestingly, the state of the node representing the Interleukin 2 (IL2) cytokine activation illustrates the differences of action of these receptors. Activation of the IL2 gene depends mainly on the activation of three transcription factors: the Nuclear Factor of Activated T cells (NFAT), the AP1 complex, and the Nuclear factor NF-kappa-B (NF-*κ*B) (Smith-Garvin et al., 2009). When NFAT and AP1 are both active, they form a complex and together bind a regulatory region of the IL2 gene. In absence of AP1, NFAT induces a different program leading to cellular anergy (Macian, 2005; Smith-Garvin et al., 2009): activation of Diacylglycerol Kinase (DGK) prevents DAG-mediated activation of RasGRP1, which regulates the threshold for T cell activation (Roose et al., 2007; Das et al., 2009).

Our comparative propagation analysis reveals that while the activation of the CTLA4 receptor leads to a general inactivation of the three transcription factors regulating IL2 production, activation of the PD-1 receptor leads only to the inactivation of NF-*κ*B and FOS (a member of the AP1 complex), thereby preventing the formation of the NFAT/AP1 complex, but enabling the activation of DGK. This observation is consistent with the proposal to target DGK isoforms as a complement of checkpoint immunotherapy (Riese et al., 2016; Jung et al., 2018).

As a next step, new co-inhibitory receptors recently under study, such as the Hepatitis A virus cellular receptor 2 (also known as TIM3) or the Lymphocyte activation gene 3 protein (LAG-3) (Anderson et al., 2016), could be easily added to the model described here, provided sufficient information could be gathered regarding their interacting partners. Applying propagation analysis in this context would be greatly insightful for future therapy developments.

## AUTHOR CONTRIBUTIONS

CH developed the T cell signalling molecular map and model under the supervision of MTC and DT. CH and AN implemented the computational methods and applied them to the T cell model under the supervision of AN and DT. All co-authors contributed to the redaction of the manuscript and endorse its content.

## ACKNOWLEDGMENTS

We warmly thank Bernard Malissen, Romain Roncagalli and Guillaume Voisinne, for their thoughtful guidance during the construction of the T cell signalling model.

## SUPPLEMENTAL DATA

Supplementary Figure 1: Molecular map in good resolution or vectorial format Caption: This molecular map describes biological entities and reactions implicated in the process leading to activation of CD4+ T cells in humans/mouse. The cytoplasmic membrane and attached receptor proteins are placed at the top, while the cell nucleus is located at the bottom. Reactions represent current knowledge of various pathways related to cytoskeletal remodelling, calcium fluxes, metabolism, cell cycle or IL2 production, and are encoded using the CellDesigner software (Funahashi et al., 2008) (the CellDesigner file is provided as Supplementary File 1).

Supplementary File 1: Molecular map in CellDesigner/XML format (Funahashi et al., 2008).

## DATA AVAILABILITY STATEMENT

In addition, the original model (GINsim format – zginml) can be downloaded from the GINsim website (http://ginsim.org/model/tcell-checkpoint-inhibitors-tcla4-pd1), together with previews of the two interactive notebooks enabling the reproduction of all the analyses and results reported in this study, using the tools integrated in the most recent CoLoMoTo Docker image (https://github.com/colomoto/colomoto-docker). A SBML-qual export of the logical model is also available on the GINsim model repository page, which will be further deposited into the database BioModels (https://www.ebi.ac.uk/biomodels/) upon publication.

